# High Throughput Sequencing technologies complemented by grower’s perception highlight the impact of tomato virome in diversified vegetable farms

**DOI:** 10.1101/2023.01.12.523758

**Authors:** Coline Temple, Arnaud G. Blouin, Sophie Tindale, Stephan Steyer, Kevin Marechal, Sebastien Massart

## Abstract

The number of small-scale diversified vegetable growers in industrialized countries has risen sharply over the last ten years. The risks associated with plant viruses in these systems have been barely studied in Europe, yet dramatic virus emergence events, such as tomato brown fruit rugose virus, sometimes occur. We developed a methodology that aimed to understand better the implications related to viruses for tomato production in Belgian’s vegetable farms by comparing growers’ perception of the presence of viral symptoms (visual inspection) with non targeting detection of nearly all viruses present in the plants by high throughput sequencing technologies (HTS). Virus presence and impact were interpreted considering the farm’s typology and cultural practices, the grower’s professional profiles, and visual inspection of plant-viral-like symptoms. Overall, The data indicated that most growers have limited understanding of tomato viruses and are not concerned about them. Field observations were correlated to this perception as the prevalence of symptomatic plants was usually lower than 1%. However, important and potentially emergent viruses, mainly transmitted by insects, were detected in several farms. Noteworthy, the presence of these viruses was correlated with the number of plant species grown per site (diversity) but not with a higher awareness of the growers regarding plant viral diseases or a higher number of symptomatic plants. In addition, both HTS and perception analysis underlined the rising incidence and importance of an emergent virus: Physostegia chlorotic mottle virus. Overall, the original methodology developed here, combining social science with HTS technologies, could be applied to other crops in other systems to identify emergent risks associated with plant viruses and can highlight the communication needed toward growers to mitigate epidemics.

## 1 Introduction

Tomato (*Solanum lycopersicum L.*) is one of the most popular and valuable cultivated vegetables grown worldwide with a gross production of 102.6 billion US dollars and yield estimated at 186.8 million tons (MT) in 2020 (FAOSTAT, 2020). Tomatoes are grown in diverse production systems (open fields, under plastic tunnels, hydroponics, high-tech greenhouses) for the fresh market or food industry. It is Europe’s main produced vegetable with 18 MT in 2021 (Eurostat, 2021). During this year, the major part of the supply was in Italy (6,6MT) and Spain (4,7 MT), where production occurs in open fields and tunnels, mainly for processing and export. In northern Europe, Poland (1,1 MT), Nederlands (0,9MT), and Belgium (0,3MT) are the largest tomato producers, specializing in the production of fresh edible tomatoes in high-tech greenhouses (Eurostat, 2021).

Tomatoes are also grown by small-scale growers and in gardens for local consumption (Benton Jones, 2007). In industrialized countries where small-scale growers almost disappeared during the green revolution, these production systems represent a niche market which is recently expanding (Morel and Leger, 2016, Laforge et al., 2018, Dumont et al., 2020). These small-scale growers often promote human values and ecosystem welfare rather than profit maximization (Morel and Leger, 2016). Combining multiple logics and aspirations is indeed typical of agroecology-inspired growers (Plateau et al., 2021). Regarding farming practices specifically, most of these growers aim to sustainably produce an extensive range of vegetables on soil, leaning on eco-systemic services, crop diversification, and rotations. Studies have shown that these systems have many advantages over conventional agriculture, especially for the environment and workers, as it reduces chemical and polluting intrant. In addition, these systems are supposed to have better resilience to climate change and plant diseases (Kremen et al., 2012, Mori et al., 2013, King and lively, 2012). It has also been shown that multi-cropping and crop rotations increase yields in both organic and conventional cropping systems (Ponisio et al., 2015), encouraging the need for research on these agricultural practices to improve the productivity of sustainable agriculture methods.

In Belgium, there are two distinct sectors of tomato production. The most significant part of tomato production is dedicated to export and mass retailing. It is cultivated mainly in the northern part of the country (Flanders) by specialized tomato growers under high-tech greenhouses. On the other hand, a minor part of the production is achieved by small-scale growers producing tomatoes amongst other vegetables for local consumption. The number of these small-scale (< 2ha) growers has risen sharply over the last ten years in the southern part of Belgium, Wallonia (Dumont et al., 2020). A sociologic survey underlined that ethical and sociological factors were considered in their decision processes and that many of these growers are not from the agricultural sector (Dumont 2017). Most of these growers aimed to produce tomatoes on soil under tunnels or greenhouses and alternated with other vegetables over a year (Dumont 2017). Tomatoes are sensitive crop: in all growing systems, the presence of pests and diseases (fungi, bacteria, viruses…), can jeopardize tomato crop, leading to important yield losses (Blancard, 2009). The characteristics of each pathosystem (the subset of an ecosystem in which the components include a host organism and an associated pathogen or parasite) determine specific strategies for plant pathogen control (Aranda, Freitas-Astúa, 2017). Thus, viral outbreaks are often related to unknown emerging diseases, for which diagnosis is the first step in disease management (Hanssen et al., 2010, Rubio et al., 2020).

Viral diseases represent nearly half of the emerging plant diseases (Anderson et al., 2004) and tomato is the plant for which the most viruses are recorded (312 viral species in 2021, Rivarez et al., 2021). Plant viruses can be spread through insects, seeds, plant-to-plant contact, fungal spores, and other means.

Many environmental factors drive the emergence of plant viruses and their outbreaks by altering interactions between viruses, hosts, and vectors, if any. For plant viruses, climate change and human activity, such as agriculture and trade, are the main factors influencing the outcome of these interactions (Jones et al., 2009, Elena et al., 2014). Elena et al., (2014) decoupled the emergence of new viruses into three phases. The first one requires a virus to jump from a host (“reservoir”) to the same host in another ecological environment or to a new host (“spillover”). This phase can be facilitated by introducing a new plant species or vector into a geographical area where they were not present. For example, the ecological behavior of insects, the main vectors of plant viruses, may be disturbed by climate change, which can result in new virus outbreaks (Jones et al., 2016). Trade and exchanges of plants between different geographical area (e.g. from their crop domestication center to a new area) has also been found to be the origin of virus emergence and epidemic (Anderson et al., 2004, Jones et al., 2009). The second phase involves the adaptation of the virus in its new host/environment in which it develops the ability to be transmitted independently from the reservoir. The last phase is characterized by optimizing the virus transmission in this new host/environment and establishing the pathogen in the host population (Elena et al., 2014). The last two phases can be facilitated by the intensification of agriculture (i.e. monoculture) as high densities of host species facilitate pathogen establishment and transmission within the new host (Keesing et al., 2010). In addition, it is supposed that ecosystem simplification, which is suitable for a high rate of horizontal transmission (Keesing et al., 2006) would favor the evolution of highly pathogenic viruses isolates for viruses with narrow host range (Fraile et García-Arenal 2016).

Although many viruses are known to infect tomatoes, all the interactions between the different actors of the pathosystem (vector, host, virus) must occur in a favorable environment for a virus to lead to an epidemic and consequences on the production. For annual crops, viruses can be present without causing problems if their horizontal transmission is inefficient, resulting in a low prevalence in the crop over cultural season. Therefore, the presence or absence of a virus in a given environment does not necessarily reflect the health of a field and is not always equivalent to the disease impact (“viral disease risk”) but is the first step in understanding and predicting possible risks (MacDiarmid et al., 2013). The development of high throughput sequencing (HTS) technologies significantly improved the detection of new and potentially emergent viruses in the last decade (Massart et al., 2014). For example, it helped to carry out surveillance studies for tomatoes without a priori (Xu et al., 2017, Desbiez et al., 2020, Rivarez et al., 2021, Vučurović et al., 2021), to identify emergent new viruses such as tomato brown rugose fruit virus (ToBRFV) (Salem et al., 2016), Physostegia chlorotic mottle virus (PhCMoV, Menzel et al., 2018) and to study their evolution and epidemiology (Lefeuvre et al., 2019).

Of these emerging viral diseases, ToBRFV, which belongs to the *Tombamovirus* genus, has recently received the most attention from European scientists, policymakers and regulators and has sparked waves of regulatory action (Oladokun et al., 2019). ToBRFV is recommended to be regulated as a quarantine pest by EPPO (https://www.eppo.int/ACTIVITIES/plant_quarantine/A2_list). This phenomenon is due to the association of ToBRFV with severe yield losses on tomato and pepper, combined with very high contagiousness (transmission by contact: tools, hands, clothes…) and with the difficulty of sanitation (it can remain active in the environment for months) (Oladokun et al., 2019, Zhang et al., 2022).

PhCMoV also raised concerns as it is associated with extreme symptoms in tomato fruits. However, since it was only detected at a low prevalence in the field so far, the threat of the virus was supposed to be lower. This virus has a vast host range spanning across nine families and infecting crops (eggplant, cucumber, crosne), weeds (galinsoge) and ornementals (helleborus, etc.) (Temple et al., 2021). It is likely transmitted by leafhoppers such as its close relative *Alphanuclorhabdovirus*, eggplant mottled dwarf virus (EMDV) and potato yellow dwarf virus (PYDV) (Babaie et al., 2003, Black, 1942).

Since viruses cannot be cured, their control mainly relies on the use of resistant varieties or limitation of their transmission, which can be either horizontal or/and vertical (seeds) and depends on the biological properties of each virus (Hull et al., 2014, Nicaise et al., 2014). Vega et al., (2019) propose to classify pathogens based on their dispersal and survival strategies, regardless of the taxonomic group to which they belong. This classification facilitates the interpretation of the occurrence of a viral disease in response to cultural practices. For example, insecticide use will limit the presence of vector-transmitted viruses but not of seed-transmitted viruses. Furthermore, viruses can easily be mistaken with abiotic stresses, such as nutrient deficiencies (Jones et al., 2014). In addition, plant virus testing can represent a limitation for the growers and stakeholders because of the cost and accessibility. Hence, growers mainly rely on their observations and knowledge to control virus infection in the field. In this context, it is crucial to determine virus perception by growers to understand the global virus-associated risks because their actions can challenge the spread of a viral disease (Murray-Watson et al., 2022). In addition, they are the first to observe the crops and to be conscious of their loss. Still, their perception of virus infection can sometimes be disconnected from reality, leading to inappropriate practices (e.g., using fungicides to control insect-transmitted viruses) (Schreinemachers et al., 2015). Growers’ perceptions and actions depend on several factors, including their knowledge of the disease, their virus-related experience (e.g., if they have observed the virus before) and their production systems per se (e.g., if their activity depends only on one crop or on several ones). Furthermore, the growers’ actions are constrained by their financial means. In connection with the chosen production systems, the aspirations can also influence how growers deal with viruses: some producers may value growing vegetables more sustainably (which relies on ecosystem services and promotes ecosystem welfare) than maximizing their profit (Morel et al., 2016). Therefore, they would tend to display different cultural practices than “conventional growers”, which may play a role in virus presence and disease transmission. For example, growers who emphasize ecosystem welfare can be more reluctant to use insecticides to control the spread of insect-transmitted viruses as it kill other non-targeted insects that might be important for other ecological functions (pollisation, auxiliaries…). Another example is that they might be more likely to grow various tomato varieties, including old varieties that are not resistant to certain viruses such as tomato mosaic virus (Hansen et al., 2010) or, re-use their own seeds, which can promote the spread of seed-transmitted viruses.

Considering the importance of studying plant viruses (emergent or not) before they become a problem, the recent threat of ToBRFV in Belgium, and the crisis context (climate change, trade, sustainable agricultural challenges), this study aims to evaluate and compare the diversity of viruses in tomatoes grown on soil in diversified production systems with the associated risk perception of growers. Therefore, the first objective is to identify and understand the potential risks of viruses (“viral disease risk”) in these production systems in Wallonia, Belgium.

Duong et al., 2019 emphasize that although biosecurity is the second largest risk mentioned by producers, there is a lack of research on socio-economic factors that explains risk perceptions, especially those that influence risk perceptions related to biosecurity. In addition, cultural practices’ role in the viral presence and disease risk is critical to understand plant virus epidemiology (Jones, 2014), especially in sustainable agriculture where options for handling viral diseases are restricted. Therefore, the secondary objective is to interpret the results considering the farm’s typology and cultural practices, the grower’s professional profiles, and the visual inspection of plant-viral-like symptoms and to potentially evaluate what would drive the presence of viruses and their impact. In this study, HTS will be used to assess the presence of viruses without a priori. Growers’ perceptions will be compared to the presence of viruses and to the observations on the field to understand better the disease risk associated with these viruses within the different farms.

## 2 Material and methods

### 2.1 Study design

To better understand the implications of plant viruses for tomato production in Walloon vegetable farms, a three-tiered survey was designed: 1) Interviews with the owner or manager of the tomato production, 2) Field observations, and 3) Analysis of the tomato virome through HTS technologies.

In 2020, a pilot survey was carried out with five growers and three members of the Interprofessional Center of vegetable growers (CIM, Regional extension services supervising vegetable production in Wallonia) to test and improve the study design and to homogenize the questionnaire. Members of the CIM mentioned that they barely encountered outbreaks due to viral diseases on vegetable crops, including tomatoes: “most of the time, there are few virus-infected plants here and there, but viral epidemics are uncommon”. For them, tomatoes’ significant problems are related to cryptogamic diseases.

A year after this pilot survey, the study was conducted with a standardized questionnaire with 21 tomato-growers in the province of Namur and Walloon Brabant at the end of the growing season (from August 18^th^ to October 1^st^ 2021) because the prevalence of viral diseases is usually highest at this time since the viral infection has been building up throughout the season.

### 2.2 Semi-structured interviews

#### 2.2.1 Data collection

Growers’ contact details were collected through the CIM, informal growers’ network and by word of mouth.

Interviews were conducted face to face with the grower, informing the survey’s objective before the visit. During the exchanges, notes were taken, and interviews were audio recorded with the grower’s approval. The questionnaire had two main objectives:

1) to describe the typology of the farms, grower’s profiles, and cultural practices of tomato growing and, 2) to evaluate the perception of growers regarding Tomato viral diseases

First, information about the farm (farm age, area, number of vegetable species grown…), tomato culture (number of plants and varieties grown, seedling origin…) and professional background of the interviewed person (number of years in the field, family in the sector…) were collected. The questionnaire is described in Supplementary data, and the answers for each farm are presented in Supplementary table 1. To obtain a global view of the answers, median, average, min and max values were calculated for the quantitative data and the sum for the binary data.

The second part of the questionnaire evaluated how growers perceived tomato viral diseases and which control measures were applied or envisioned. A mix of several “open-ended” questions encouraging discussion and closed questions were asked in a specific order (Figure 1).

**Figure 1.**
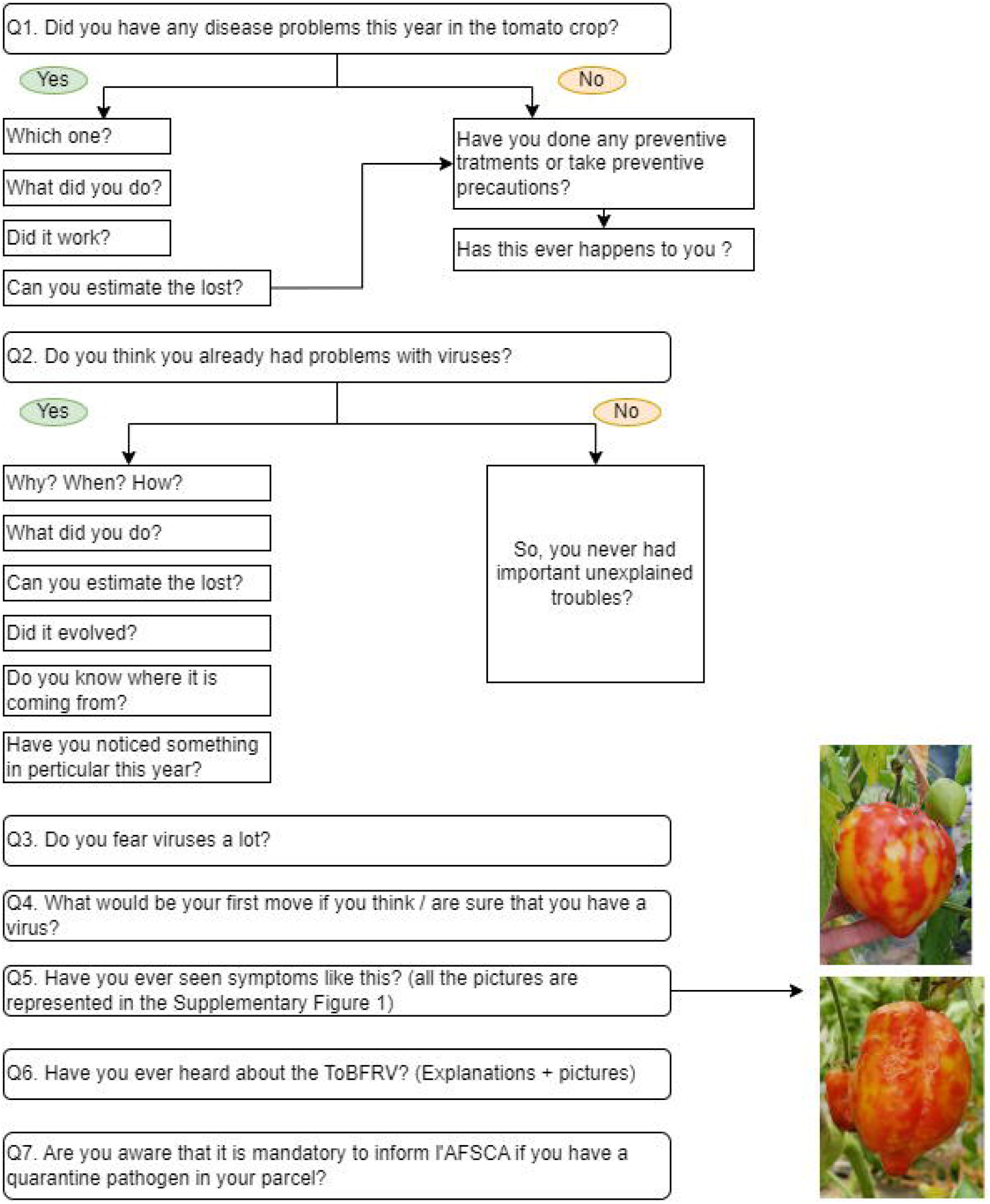
Questions related to virus’ perception

At question Q5, pictures of viral symptoms induced by PhCMoV on different host plants were shown to assess if the growers recognized the symptoms (Supplementary Figure 1). Since viral symptoms are difficult to notice and sometimes resemble other stress, this allowed the respondents to validate or correct their answers to Q2, “Do you think you have ever had virus problems”?

PhCMoV symptoms were chosen because they are severe and can be easily identified in tomato fruits. They can also be mistaken with other essential plant viruses known to be present in Belgium, such as ToBRFV or PePMV (Temple et al., 2021, Hanssen et al., 2009, EPPO Bulletin, 2020). In addition, it was the most frequently detected virus-causing symptom during the pilot survey in 2020.

In Q6, whether growers’ were aware of ToBRFV, was investigated as this virus was recently widely publicized by different stakeholders involved in the tomato production chain.

At the end of the interview, information on the biology of these two viruses (PhCMoV & ToBRFV), which require different control measures, were given to the growers.

#### 2.2.2 Data interpretation

The second part of the questionnaire mobilized the perception and worries of the growers and allowed them to express themselves spontaneously. Interviews were first transcribed word by word, and answers with the same meaning were grouped manually. For instance, concerning Q2, the growers all expressed their answers differently. Their responses were classified into four categories: those who thought they already had viruses, those who did not say “yes” clearly but who “did not rule out the possibility”, those who “did not know”, and those who “did not think” they already had viruses.

### 2.3 Observations and sampling

Before sampling, each grower was explicitly told that if a quarantine virus (eg. ToBRFV) was detected, it was mandatory to notify Belgian’s NPPO (Federal Agency for the Safety of Food chain). After that and prior to sample collection, oral consent for sampling was given.

In each farm, 100 asymptomatic tomato leaves were systemically collected in a W shape. When there were several tunnels on a farm, an equal number of plants was sampled per tunnel to reach 100 plants per farm. The tomato plants that showed viral-like symptoms (fruits: deformations, anomalies of coloration; leaves: vein clearing, deformation, mosaic; plant: size) were pictured, counted, and collected in a separate bag.

Since the symptoms of PhCMoV can be easier to spot on eggplant than on Tomato, and PhCMoV was the virus causing symptoms the most detected during the pilot survey, we also looked at eggplants when present on the farm. When the symptoms of PhCMoV were noticed on eggplants or tomatoes, the number of symptomatic plants was recorded, and at least three different symptomatic plants per farm were collected.

In Belgium, the season 2021 was not optimal for outdoor tomato production due to very wet, dark, and cool weather (completed with storms and floods). These conditions favoured the development of fungal diseases, asphyxiated the root systems, and slowed down the ripening of the fruit. Consequently, some growers removed a part of the planting before the visit. These growers were listed (Supplementary Table 1).

### 2.4 Virus analysis

#### 2.4.1 HTS

After the collection, 100 mg of fresh tomato asymptomatic leaves from the same farm (i.e. 10 g for 100 plants) were pooled in a filter bag and stored at −80°C. The symptomatic plants were also pooled per farm, and the weight of material per plant varied according to the number of plants in total (5 g in total). After that, the samples were analyzed for viruses using a virion-associated nucleic acids enrichment protocol (VANA) before HTS on Illumina. The VANA protocol and library preparation used for the samples followed the method described by Maclot *et al*., (2021) adapted from Filloux *et al*., (2015).

In brief, 5 or 10 g of tissue were ground respectively in 25 or 50 mL of a cold Hanks’ buffered salt solution (HBSS, composed of 0.137 M NaCl, 5.4 mM KCl, 0.25 mM Na_2_HPO_4_, 0.07 g glucose, 0.44 mM KH_2_PO_4_, 1.3 mM CaCl_2_, 1.0 mM MgSO_4_, 4.2 mM NaHCO_3_) using a tissue homogenizer. In a 50 mL falcon tube, the clarification was obtained from a centrifugation run of 10,000 g for 10 min at 4°C. Supernatant was then filtered through a 0.45 *μ*m sterile syringe filter and 10,4 ml of supernatant were put into an ultracentrifuge tube (Beckman Ultra clear 14 mL tubes (#344085)). In each tube, 0.1 mL (1:100) of a clarified banana sample infected with banana bract mosaic virus (BBrMV) was added as an internal positive control to evaluate the analytical sensitivity of the test (Massart et al., 2022). Then, a sucrose cushion, made of 1 mL of 30% sucrose in 0.2 M potassium phosphate at pH 7.0, was deposited at the bottom of the tube. Extract was then centrifuged at 40 000 rpm for 2 hours at 4°C using the 50Ti rotor (Beckman). A protocole previsouly used in the laboratory and described in Maclot et al. (2021) was used for the library preparation.

PCR products were pooled by 6 to 12 according to the linkers and cleaned using the Nucleospin Gel and PCR clean up (Macherey-Nagel). Samples containing asymptomatic plants were pooled separately from those containing symptomatic leaves to limit potential cross-contaminations of highly concentrated viruses in symptomatic pools. A positive external alien control containing infected beans with Endornavirus was processed simultaneously as the asymptomatic samples to monitor potential cross-contaminations as recommended in Massart et al., (2022).

Illumina library was prepared at GIGA Genomics (University of Liege, Belgium) using NEBNext Ultra II DNA library prep kit (New England BioLabs, US) and libraries were sequenced on the Illumina NextSeq500 sequencer for the generation 10 M of paired-end reads (2 x 150 base pair) per library. Resulting sequence reads were first demultiplexed according to the linker and trimmed from the adaptor, then quality trimming, pairing, and merging were performed using the Geneious R11 software platform (https://www.geneious.com) before de novo assembly with (RNA) SPAdes assembler 3.10.0 (Bankevich et al., 2012). Next, contigs were compared using tBlastx against a database of viruses and viroids sequences downloaded from NCBI in November 2021 (RefSeq virus database). Then, reads were mapped on the closest reference sequences using geneious parameters Medium-Low Sensitivity/ Fast. The presence of viruses was considered positive when the coverage (% of reference genome) was superior to 50% for most viruses and when the number of mapped reads on the closest genome reference was > 90. On another hand, the presence of tombusviruses and alphanecroviruses was assessed at a different threshold (12% of ref seq) since the number of reads which map on reference genomes, were very low compared to other viruses, such as the internal control BBrMV. This threshold was determined after a manual expertise assessment of the difference between contamination and low concentration of each virus (Rong et al., 2022).

#### 2.4.2 RT-PCR for PhCMoV detection in eggplant

Eggplant leaves showing symptoms of PhCMoV were subjected to RNA extraction and RT-PCR for testing the presence of PhCMoV. RNA extracted followed the method described by Onate-Sanchez and Vicente-Carbajosa (2008). Then, the extracts were reverse transcribed using random hexamers and Tetro RT enzyme (Bioline). The obtained cDNA was amplified with the MangoTaq™ DNA Polymerase and the primers described by Gaafar et al., (2018). Thermal cycling corresponded to: 94°C for 1 min, 35 cycles at 94°C for 15 s, 60°C for 20 s and 72°C for 45 s, with a final 72°C extension for 3 min. Amplicons products were analyzed by electrophoresis on a 1% agarose gel in Tris-acetate-EDTA (TAE) buffer stained with GelRed^®^ Nucleic Acid Gel Stain (Biotium) and visualized under UV light.

#### 2.4.3 RT-PCR and sanger sequencing for confirmation of challenging HTS results

To confirm the detection of strawberry latent ringspot virus (SLRV) and carnation Italian ringspot virus (CIRV), RNA from the original 100 frozen leaves of the positive sample were re-extracted in pools of 25 with the Spectrum plant total RNA kit (Sigma-Aldrich) and tested by RT-PCR using the Titan One Tube RT-PCR kit. For the detection of SLRV, the primers SLRSV1/2 described by Bertolini et al., were used (2003) and for the detection of CIRV, primers were designed based on the consensus sequence using Geneious designing primer tool: CIRV-F: “CGTGGCAGTTACCAGACAGT”, CIRV-R: “CTCCATCCCAACGTTCACCA” (product length: ~1kb). Amplicons were Sanger-sequenced and the obtained sequences were aligned to the respective reference sequences using BioEdit to confirm the presence of the virus.

The status of tomato mosaic virus (ToMV) was assessed in six six samples where HTS yielded a small number of reads mapping the reference genome. The 100 frozen leaves for each of these sample and the positive sample were extracted in pools of 25 following the Spectrum plant total RNA kit (Sigma-Aldrich). RNA extracts were tested by RT-PCR using the Titan One Tube RT-PCR kit (Roche) and the primers of Li et al., 2018.

### 2.5 Data comparison

In this study, we classified the detected viruses based on their transmission mode (insect-vector, soil, seeds, fungi, unknown, Table 1a). After that, the number of sites where viruses transmitted in the same way were detected were counted and gathered in a same category (Table 1b). Further analyses were made on the category with the higher number of detections within symptomatic plants since this feature may be linked to the grower’s perception.

**Table 1.**
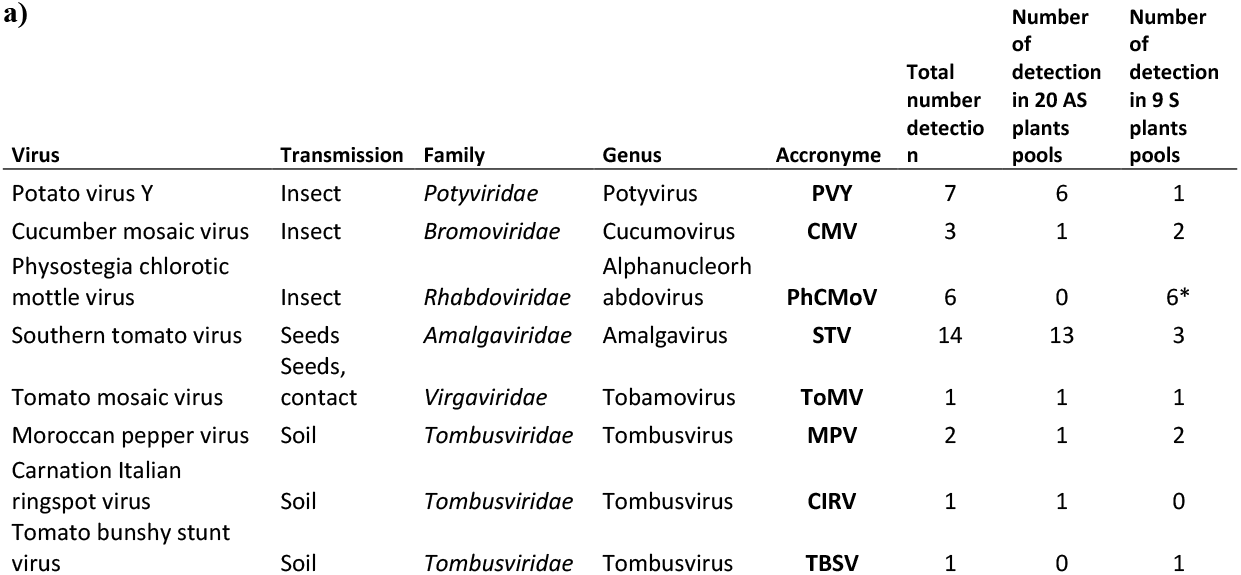
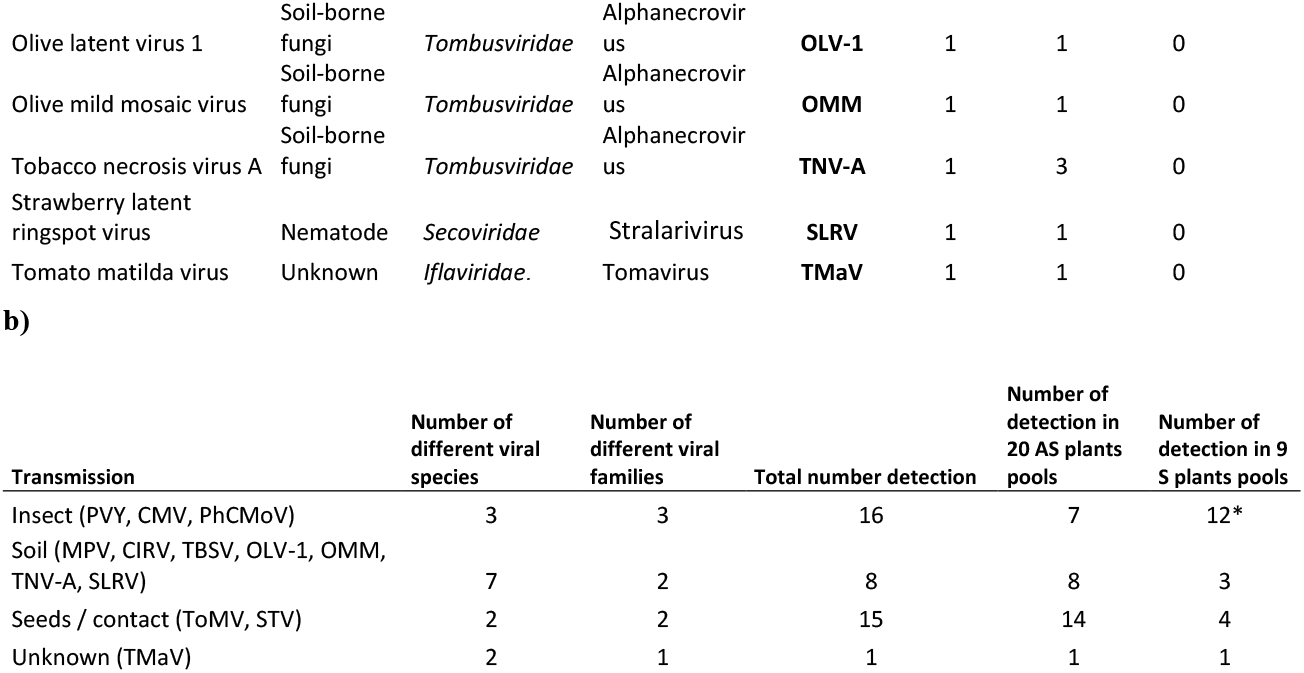
Taxonomic characteristics of detected viruses and the number of farms where they were detected. Whether the detection occurred in symptomatic (S) or asymptomatic (AS) plants pools is indicated (a). Viruses were classified according to their transmission mode (insect, soil, seeds, unknown), and the number of detections for all the viruses in each mode is indicated in b) *Three detections were only made on symptomatic eggplant by RT-PCR (Supplementary table 2)

The most prevalent transmission mode was further used to compare the site where viruses transmitted with this mode were present to the others. For that purpose, quantitative data were transformed into qualitative data based on the median number of each feature. In addition, all the data were normalized according to the number of growers with or without the viruses transmitted the same way.

WalOnMap (https://geoportail.wallonie.be/home.html) was used to determine the agricultural area where the different farms were located, and growers were grouped based on agricultural area and geographical relevance (growers situated in the adjacent area of sandy-silty and silty were grouped and the ones in the adjacent Condroz and Famenne areas as well).

In addition, to assess whether the correlations were significant, all the generated raw data were tested for correlations using Orange Mining’s Sieve Plot diagrams (Demsar et al., 2013). Finally, using the relevance scoring options, the variables that were most associated with the presence of some viruses transmitted in the same way or interesting results linked to grower’s perception were identified.

## 3 Results

### 3.1 General description of the farms, professional profile of the growers, and tomato culture

During the first step of the questionnaire, information was gathered on the general farm’s characteristics, the professional profile of the growers, on tomato cultivation practices. The detailed data are presented for each farm in Supplementary table 1.

The 21 surveyed growers sold their products locally (e.g. shop in the farm, markets, baskets, “pick your own”). The median time they have worked in vegetable farming is seven years, with only four years (median) of tomato production on the studied site (Fig. 2a). This difference is because most of them worked somewhere else before starting their own production but when they did, they have always grown tomatoes. Some growers (9/21) only worked in vegetable cultivation, while others (12/21) switched careers after having worked in other sectors (Fig. 2b). Only five respondents have relatives in the agricultural sector (Fig. 2b).

**Figure 2.**
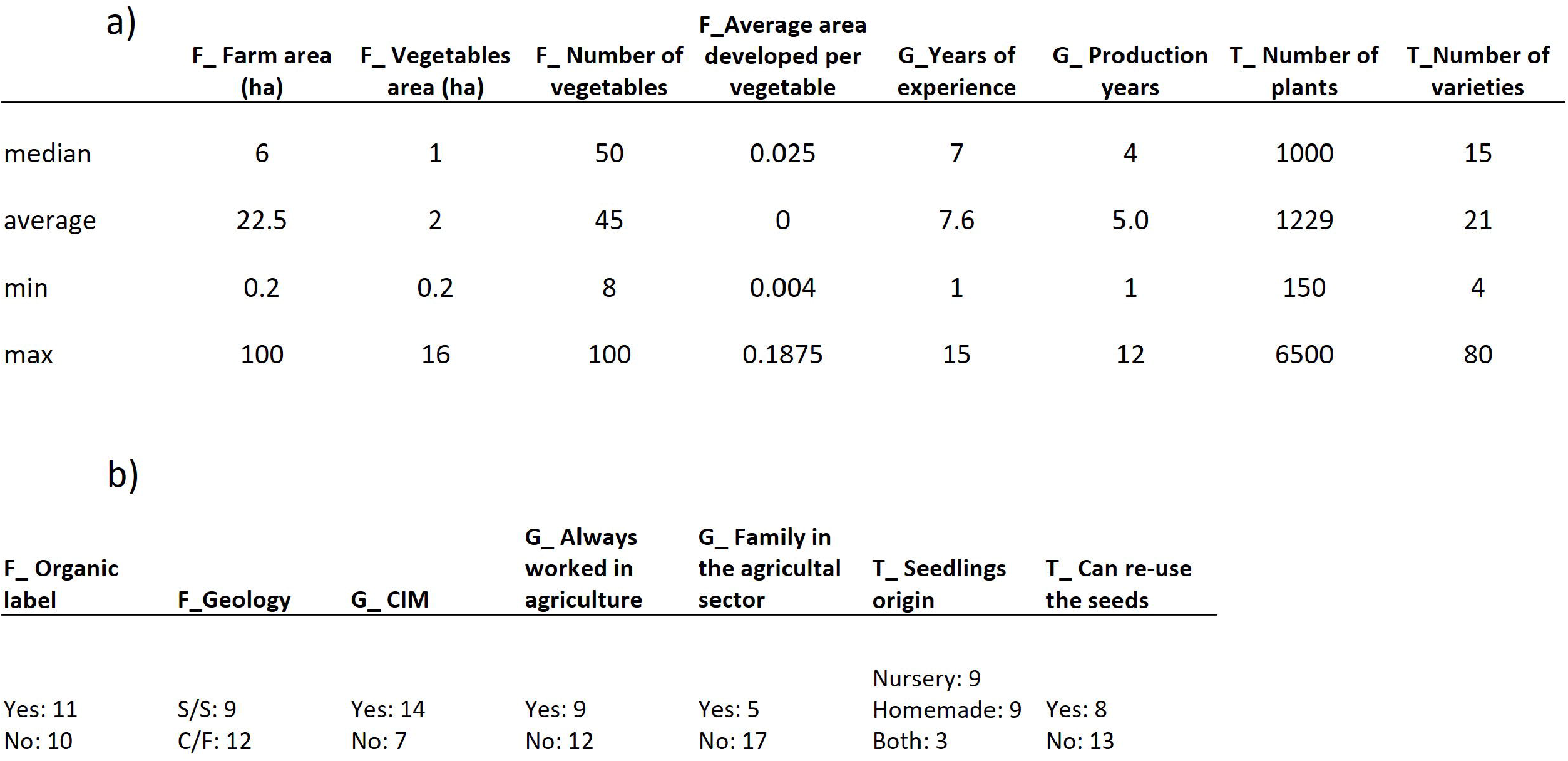
Description of the main characteristics of the farms (F); grower’s profiles (G) and, tomato cultural practices (T) of the respondents (n=21) a) Median, Average, Min and Max values of quantitative data b) Qualitatives data, S/S = silty and sandy-silty, C/F = condroz and famenne area

Regarding growing systems, half of the respondents grew vegetables under the organic label or were in the process of obtaining it. However, among the other half, most growers followed organic or agroecological farming practices without certification. Moreover, some of them explained that they don’t need the organic label while following the practices because they are “close to their consumers” (notably in the case of pick-your-own farms). Therefore, it is challenging to mobilize this feature in the analyses.

Most of the surveyed growers produced many different vegetable species per year (median = 50) on a small surface area (median = 1.1 ha) (Fig. 2a). The different crops (root vegetables, fruit vegetables, leafy green, cruciferous, marrow, aromatics…) alternate on the same piece of land throughout the year. Only two growers stood out from the others by growing vegetables on a larger surface (INX-29: 4.5 ha and INX-37: 16 ha) (Supplementary table 1).

Regarding tomato production, the median number of tomato plants grown per year and farm was 1,000, with a median of 15 different cultivars. The growers bought their plants from nurseries or made seedlings from commercial or home-made seeds from the previous year (Fig. 2b).

### 3.2 General description of the grower perception, observations of viral-like symptoms and virus detected in the 21 farms

#### 3.2.1 Grower perception

During the interview, 20 out of 21 growers responded having faced cryptogamic diseases during the current year (Fig. 3a - Q1). At that stage, a single grower (INX-40) mentioned encountering problems with viruses.

**Figure 3.**
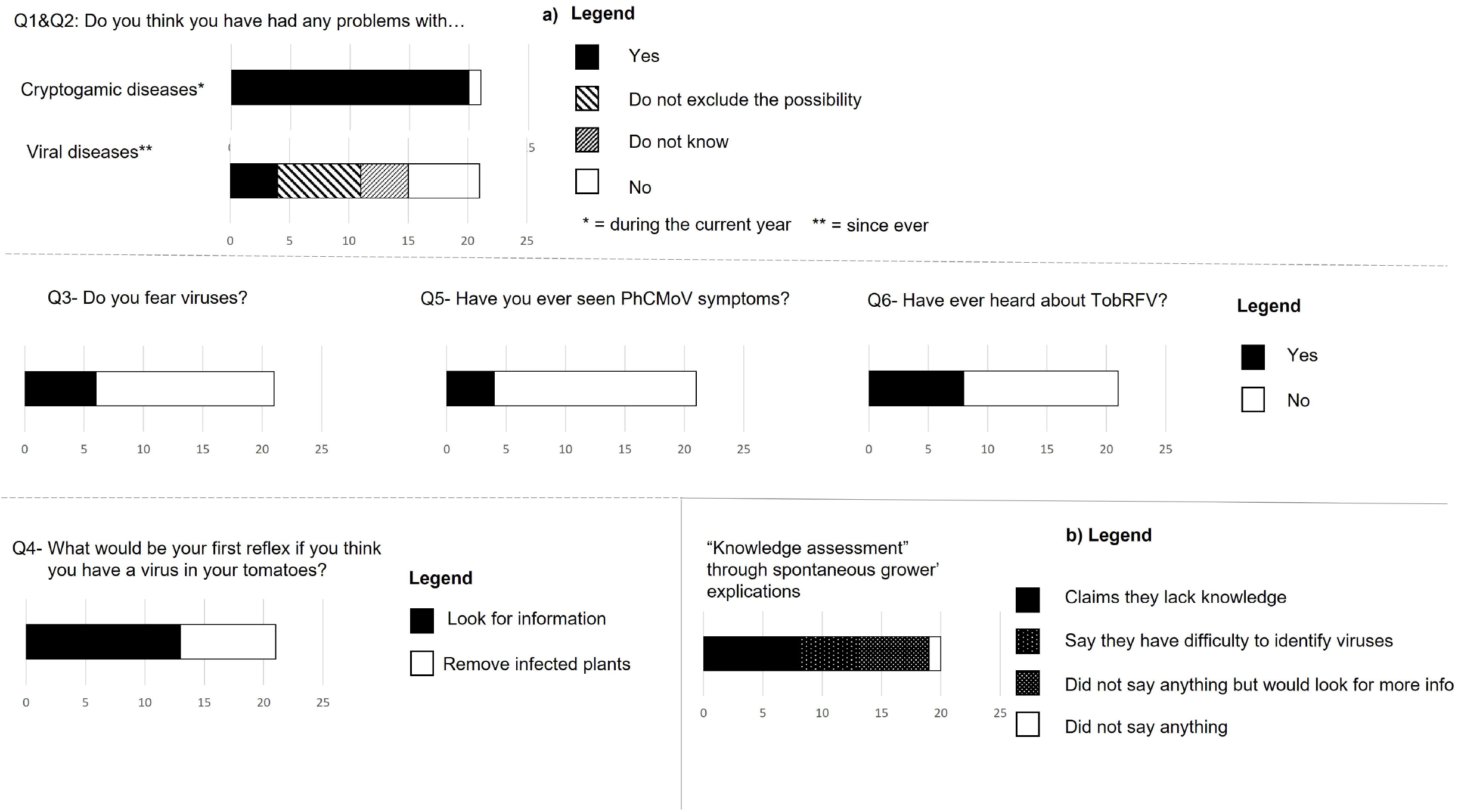
Grower perception (n=21) regarding tomatoes diseases focus on viral diseases. a) answers to perception’s study and b) interpretation of the interviews related to virus knowledge

In the second question specific to viral diseases, only four growers responded that they had already faced virus infection, but it had never been problematic (except for the grower INX-40). Seven other growers did not rule out the possibility of having viruses but were unsure. The rest of the growers “did not know” or “did not think” they ever had tomato infected by viruses (Fig. 3a - Q2). Interestingly, no growers highlighted that they already had terrible unexplained troubles with tomato production at Q2. In addition, most growers (15/21) responded that they did not fear viruses (Fig. 3a - Q3).

Subsequently, the interviews revealed that many growers addressed their level of knowledge about viruses by themselves. Many respondents naturally mentioned that they “didn’t know about viruses”, that they were “unaware of them”, or that they “didn’t know how to recognize them”. The only grower that demonstrated his knowledge about plant virus was the one with the most significant tomato production (INX-37). The lack of knowledge was exposed by the question Q6 where less than half of the growers were aware of the potential danger posed by ToBRFV. They also mentioned that they did not know the list of quarantine pathogens and how to recognize them. In contrast, the growers seemed aware of fungal diseases as they all mentioned one disease name in Q1, and none highlighted their lack of knowledge about fungal diseases.

#### 3.2.2 Observations

In the parcels, 122 viral-like symptomatic tomato plants were collected in nine farms. In most cases, the proportion of symptomatic tomato plants was lower than 1% (Supplementary table 1).

Only one exception was noticed with the grower INX-40, who was aware of his viral problem (Q1, Q2). In this farm, 40 tomato plants/300 (13%) showed strong viral symptoms like the ones associated with PhCMoV (Supplementary Fig. 3). In this farm, there were three tunnels, and one of them was particularly impacted with 38/85 tomato plants showing these typical symptoms. In this tunnel, other plants were grown (cucumber, capsicum, mint, strawberry…), and the grower mentioned that he saw the symptoms related to PhCMoV in cucumber early in the season. In addition, three eggplants near the greenhouse also showed the symptoms of PhCMoV and were tested positive.

In total, 30 eggplants showing PhCMoV symptoms (vein clearing on the leaves) were observed across five farms (including two farms where symptomatic tomatoes were observed, Supplementary table 2). The maximum number of symptomatic eggplants per farm was 11 (Supplementary table 2).

#### 3.2.3 Virus detection

First, the spike BBrMV was detected in all the asymptomatic pools of 100 plants with a number of reads mapped on the reference genome NC_009745 ranging between 5 and 99. The external alien control (endornavirus) which was processed at the same time as the asymptomatic samples, was also detected in the expected samples in the two pools with 27,690 and 43,744 reads mapped on the NC_038422 genome.No reads of endornavirus were detected in other samples, suggesting that the level of cross-contamination was low.

In total, 13 different viral species belonging to eight viral families were identified during this survey (Table 1a). The number of virus species detected per farm varied between 1 and 4 (Supplementary Table 2).

These viruses were classified into four categories based on their transmission “mode”: transmitted by insects, by the soil (including virus transmitted by soil-borne fungi or nematodes), by seeds and/or contact and the viruses for which the transmission is not known to date (Table 1b). In total, three viral species transmitted by insects were detected across 16 different farms, seven viral species transmitted through the soil across nine farms, two viral species transmitted through seeds and/or contact across 14 farms, and one species for which biological data on its transmission was lacking, on one farm. In the symptomatic plant pools, insect-transmitted viruses were the most prevalent, as they were detected across 12 farms out of 21 (Table 2b).

The most frequently detected virus (n=14) was southern tomato virus (STV), a persistent virus transmitted by seeds. STV was detected more frequently in asymptomatic plant pools than in symptomatic pools (Table 1).

After that, the most prevalent viruses were the ones transmitted by insects: potato virus Y (PVY), cucumber mosaic virus (CMV), and PhCMoV, which were detected in respectively seven, three and six farms (Table 1a).

HTS identified PhCMoV in three farms on symptomatic tomato showing fruit deformations and anomalies of colouration and by RT-PCR on three additional farms on eggplants showing vein clearing on the leaves (Table 1). Overall, during the study, all the tested plants that showed PhCMoV symptoms were positive. PhCMoV was only detected in plants with typical PhCMoV symptoms and not in asymptomatic plants, whereas PVY was primarily identified in asymptomatic plants. These results are in line with the literature because symptoms associated with the presence of PhCMoV in tomatoes are severe and mainly located on the fruits (Temple et al., 2021), while the ones associated with PVY are mild leaf symptoms (Jones et al., 2014). Leafhoppers transmit PhCMoV, while aphids transmit PVY. These two viruses appeared on a farm where many symptomatic plants were found, including cucumber and eggplant and where the growers complained of tomato virus disease (INX-40). The symptoms observed were very characteristic of PhCMoV (Supplementary Figure 2c), which were detected by HTS on the pool of symptomatic tomatoes and by RT-PCR in four separate symptomatic tomato plants. PhCMoV was detected in symptomatic plants exhibiting the same symptoms without PVY. PVY has never been associated with fruit symptoms in a single infection on tomato (Jones et al., 2014), so it can be assumed that in farm INX-40, the presence of PhCMoV explained most of the observed symptoms.

CMV was associated with symptoms in two symptomatic pools and in one asymptomatic pool (Supplementary Table 2a). In addition, this aphid-borne virus has the broadest host range of any known plant virus (1200 plant species) (Gallitelli 2000).

Another virus transmitted by seeds, tomato mosaic virus (ToMV, Supplementary Figure 2b), was detected in one farm on asymptomatic and symptomatic plants. ToMV is a *Tobamovirus* which have been widely studied and is also transmitted by contact (Jones et al., 2014). RT-PCR confirmed the presence of the virus in the sample where the highest number of reads was found and denied in all the other samples.

In addition, six different viral species belonging to the *Tombusvidae* family (tomato bunshy stunt virus, moroccan pepper virus, olive latent virus 1, olive mild mosaic virus, tobacco necrosis virus A, CIRV) were primarily detected in asymptomatic plants (Table 1). These viruses are transmitted through the soil, mainly by soil-borne fungi and are not considered economically significant tomato pathogens (Yamamura et al., 2005).

Finally, SLRV and tomato matilda virus (TMaV) belonging to the *Secoviridae* and *Ilflaviridae* families were detected on asymptomatic plants only. SLRV is transmitted by a nematode, and the transmission mode of TMaV is unknown; the virus was described for the first time in 2015 and did not seem to be associated with symptoms (Saqib et al., 2015).

This reports the first detection of SLRV and CIRV on tomatoes. RT-PCR and sanger sequencing was performed on the original plant samples and confirmed the presence of these viruses in tomatoes.

### 3.3 Correlations

To investigate whether insect-transmitted viruses (PhCMoV, PVY, CMV) correlated with any specific metric related to the farm’s characteristics, cultural practices, grower’s profiles or perception, the widget sieve diagram on orange mining was used in the supplementary table 1, and correlations with a p-value < 0.1 were noted. Orange mining allows testing all the correlations between different features without a priori and scores the best correlations.

Regarding cultural practices, the presence of these three viruses was correlated with the increased number of different vegetable species grown per farm (“diversity”). Furthermore, these insect-transmitted viruses were also correlated with being situated in the silty or sandy-silty area. Finally, regarding perception and actions, the presence of PhCMoV, CMV and PVY was associated with the growers who would remove the virus-infected plants as a first reflex, with the ones who do not fear viruses, and with the ones who recognized PhCMoV symptoms (Fig. 4).

**Figure 4.**
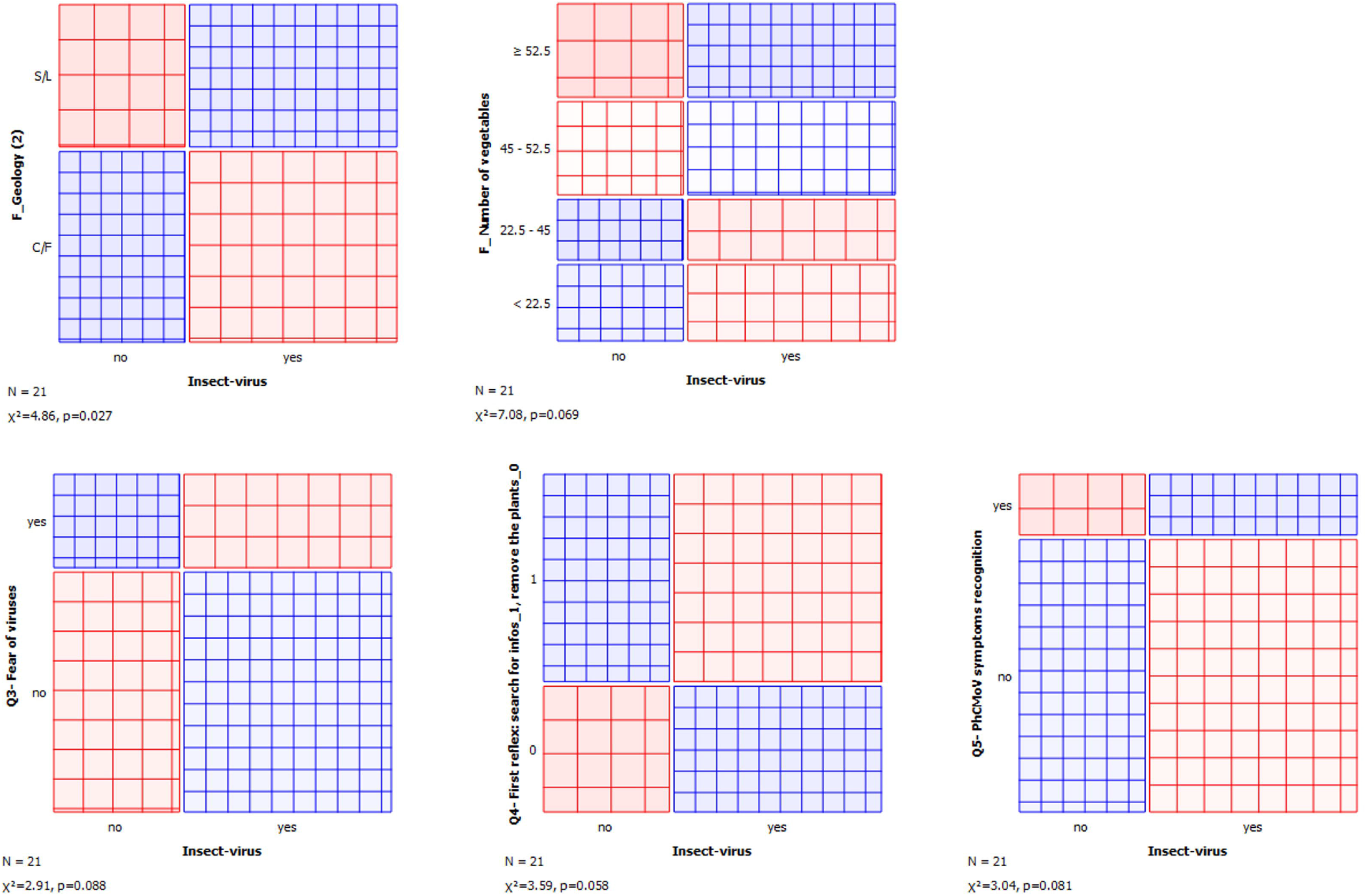
Correlations between the presence of insect-transmitted viruses and different metrics. The area of each rectangle is proportional to the expected frequency, while the observed frequency is shown by the number of squares in each rectangle. The difference between observed and expected frequency (proportional to the standard Pearson residual) appears as the density of shading, using color to indicate whether the deviation from independence is positive (blue) or negative (red). https://orangedatamining.com/widget-catalog/visualize/sievediagram/

Since PhCMoV caused remarkable symptoms in tomato fruits and represented half of the studied insect-transmitted viruses in symptomatic leaves (6 detections/12), it made sense that its presence was related to the growers who recognized PhCMoV symptoms.

In most cases, removing virus-infected plants as a first response is a good response to reduce the spread of plant viruses. Therefore, it is surprising that growers with tomatoes infected by insect-transmitted viruses mentioned they would remove the symptomatic plant if they were sure of the viral origin of the symptoms. In addition, the growers who do not fear viruses (15/21) are the ones who have the most-insect transmitted viruses. This result is also striking since insect-transmitted viruses were frequently detected in symptomatic plants. Therefore, it suggests that the growers did not notice the symptoms or that these viruses were not associated with a high prevalence in these production systems.

The number of vegetables grown per year is a feature related to the farming systems. Therefore, some other features often related to the diversity may also be correlated (albeit to a lesser extent) to the presence of viruses. In a second step, we compared if these features could be correlated with the presence of insect-transmitted viruses by simply looking at the differences between the farm where insect-transmitted viruses were found and the others. We also compared selected features of interest related to the grower’s perception or observations to evaluate how they are linked to the presence of insect-transmitted viruses (even if it is not significant). This analysis showed that, in addition to the already presented correlations, the farms where insect-transmitted viruses were detected tended to have less tomato plants, which can be related to a higher number of different plant species per year, and with the growers who produce vegetables on smaller surfaces (Fig. 5).

**Figure 5.**
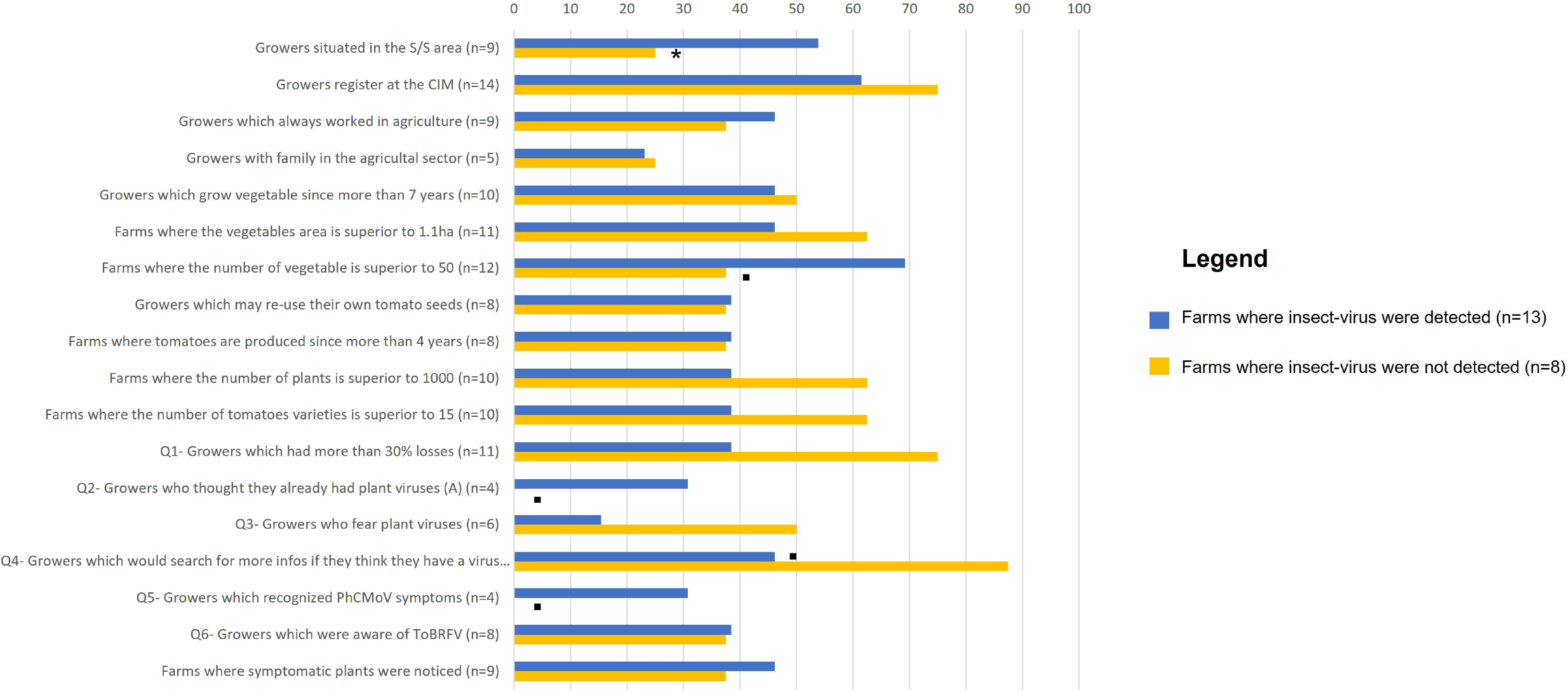
Percentage of growers which have insect-transmitted viruses (blue) or not (yellow) vs selected features. n represent the number of growers in each feature on which the analysis was based. The difference statistically highlighted with orange mining was indicated when p < 0.1 =., p < 0.05 = *

For the perception, the four growers who thought they already had virus problems and the one who recognized the symptoms of PhCMoV had insect-transmitted viruses, but interestingly, most of them did not fear viruses (Fig. 5, Supplementary table 1). These results can be related to their perception of the absence of a significant problem with viruses and highlight that the presence of PVY, CMV or PhCMoV is, in most cases, not associated with important yield losses. Also, three out of four growers who recognized the PhCMoV symptoms had PhCMoV in their farms. In addition, the growers who thought they had more than 30% losses this year due to fungal diseases on their tomato were less likely than others to have the insect-transmitted virus, suggesting that the yield losses do not result majoritarily from viral diseases in the 2021 peculiar conditions (Fig. 5). It is also interesting to notice that symptomatic plants were similarly observed in both categories of farms (ie. Those with insect-transmitted viruses and the one without). It can be explained by the fact that PVY is well represented in the insect-transmitted detected viruses, and it was detected in 6 farms on asymptomatic plants.

Since there were no significant variations (clear cut) in how growers perceived viral diseases, it was difficult to compare the perception to the farm’s characteristics and growers’ profiles, and to understand if socio-economic factors can explain plant viruses’ risk perception. Only one grower claimed that he had problems with plant viruses.

Using orange mining, it was tested whether being aware of TobRFV (n=8) or being afraid of tomato viruses (n=6) was correlated to any factors, and no significant correlations were identified.

## 4 Discussion

Overall, this study brought knowledge to growers’ perceptions regarding viruses and the occurrence of tomato viruses in Walloon’s farming systems.

The first result shows that our selection process was representative because it was aligned with the classification of Dumont (2017): most of the interviewed vegetable growers produced a large diversity of vegetables sustainably on a surface area up to 2ha.

Second, their perception of viruses was somewhat unclear: most interviewed growers found it challenging to say whether they had problems linked to viral diseases. Hence, growers usually agree that the virus is not a major problem in their tomato production. Observations of tomato plants in the field correlated with this perception, as a small number of symptomatic plants were found. However, some viruses known to affect tomato production were identified, such as ToMV, CMV and PhCMoV (Hanssen et al., 2010, Ullah et al., 2017, Mahjabeen et al., 2012, Temple et al., 2021).

Regarding the origin of the difference between perception, observations, and the presence of viruses, a first hypothesis (H1) is that the detected viruses do not cause significant problems in the agricultural system under study (low viral disease risk) and, perception and observations correlate well with reality. A second hypothesis (H2) might be that growers are unaware of the problem, and field observations are not representative because symptoms of other diseases (fungal, bacterial, abiotic stress) might mask viral symptoms. This hypothesis can be supported by the fact that some growers admitted their lack of understanding of plant viruses or difficulties for identifying them.

While H2 cannot be set aside entirely, many elements suggest that H1 better explains the difference between perception and HTS results. First, during the survey, all respondents admitted that they had a fungal disease problem and developed a control strategy for this issue. Many fungal disease problems in 2021 were observed due to weather conditions, and the results related to the general tomato diseases perception suggested that growers seemed to be aware of the disease when they were severely impacted. The second fact in favor of H1 is that the only grower who explained having virus infection on tomato had a significant virusoutbreak. The observations and laboratory analysis were very consistent with his perception: many plants with typical PhCMoV symptoms were noticed, and the presence of the virus was confirmed. Finally, in the pilot survey, the regional extension services mentioned that viruses were not a significant problem in vegetable production, and they rarely get requests on this subject. All these information suggested that viral diseases are currently not the most important diseases for tomato production in Wallonia, even if some important viruses were detected. Many reasons can explain why a virus does not necessarily cause problems in a cropping system: its transmission from plant to plant might be inefficient, and the number of infected plants, marginal (e.g. because of a low population of insect vectors or a low transmission efficiency or because the virus encounters a resistant host), the plants might get infected late, and the virus not have time to cause damages, or the temperature was not optimal for its replication into the plants (Jeger, 2020, Trebicki, 2020).

Overall, these results contrast with the situation in the province of Alméria in Spain, where viruses represent a crucial threat to tomato production (Panno et al., 2019) and novel viruses are detected at a rate of 0.9 / year (Velasco et al., 2020). Consequently, the growers in Alméria recognize the significant risk that viruses pose (Velasco et al., 2020). It shows that growers’ perception is also correlated to viral disease risk. Agricultural systems and tomato culture history may explain the difference. For example, in Alméria, tomatoes are grown intensively under plastic greenhouses at a very high density and over 8,423 ha in 2021 (Análisis de la campaña hortofrutícola. Campaña 2020/2021). There is almost no space between the parcels which can facilitate the spread of viruses between fields since there are almost no barriers between them, in contrast to the diversified farming system analyzed here generally isolated in the countryside.

In Nordic European countries, a strong interest was recently dedicated to ToBRFV. Most tomatoes produced in the north of Europe are grown irrespective of seasonality (under high-tech greenhouses). ToBFRV is under scrutiny by large tomato growers because of the high the risks caused by the large trade (e.g. fruit, equipment, packhouses, employees). This often leads to strict phytosanitary controls and a high level of awareness to all persons allowed to enter these glasshouses. Among the studied small-scale growers, ToBRFV was not detected, and less than half of the interviewed growers were aware of it. Most of them were also unaware that it is mandatory to report quarantine pests to the authorities. This was an interesting finding given the recent high-coverage in general and agricultural media for the virus, and it highlighted that improvement can be made on the communication between phytosanitary authorities and small-scale growers. These risks can be considered initially limited because trade is often restricted and local (geographically) for small-scale growers: their production can be considered as partly isolated from intensive systems of high-tech greenhouses. The lack of awareness of viruses might also be due to the system model many growers follow in this study, where other crops compensate low yield from a tomato crop a given year. This can lead to a potential underlying infection that could remain unnoticed for some time (especially if the growers are unaware of viruses or on how to identify the pathogen and report it). An outbreak would be very challenging to manage for a small farm as the virus could survive a long time in the environment (in a large glasshouse, a complete cleanup is costly and complex but possible). We did not find any socio-economic reasons or patterns to explain this low perception of ToBRFV, probably because of the small size of the study and the lack of specific questions about technical transfer, or access to information.

In this study, the most common viruses detected in symptomatic plants were insect-transmitted viruses. Our analysis showed that these viruses were more present in the farms where numerous different plant species were cultivated and, in the farms situated in the silty and sandy-silty areas. Growers with and without insect-transmitted viruses had the same profile: settled and working in agriculture for the same number of years. Therefore, it seems that technical conditions (diversity, location of the farm…) explained more than social context and grower’s actions, the presence of certain viruses in the Walloon context. It is challenging to state whether being diversified (numerous plant species cultivated), located in the silty and sandy-silty area, or whether a range of characteristics explain the presence of these viruses, but the different elements can be justified. Insect-transmitted viruses were detected more in the Silty and Sand-silty area. The Silty area is typically used for field crops such as cereals, sugar beet, potatoes, or flax. The soil is rich and fertile, and half of the Walloon farms specializing in horticulture are also found in the silty area. It is recognized that the intensification of agriculture increased the emergence of viral diseases (Roossinck et al., 2015, Pinel-Galzi et al., 2015), and the proximity of large-scale cultivated plants next to the study plots could serve as virus reservoirs (Bernardo et al., 2018). In the Walloon context and our study case, potatoes have been primarily grown in the silty area for centuries. This crop is the primary host of PVY; thus, the presence of potatoes might partly explain the presence of PVY in the farms situated in the silty region.

CMV, PhCMoV and PVY are insect-transmitted with a broad host plant range which can affect tomato yield (Jones et al., 2014, Temple et al., 2021). The results show that insect-transmitted viruses were more likely to be detected in the most diversified farms (which were not necessarily located in the silty/sandy silty area, data not shown).

Growing a high number of vegetable plant species could increase the number of host plants harbouring insect vectors (Knops et al., 2002, Cook-Patton et al., 2011), or the number of host plants enabled to host viruses (“amplification effect”, Keesing et al., 2006). Plant-cultivated diversity in small-scale production systems may also increase the first step of emergence: virus-host jump between wild and cultivated reservoirs as the number of potential new hosts is higher. In addition, these diversified farms have expanded in the past ten years throughout industrialized countries (including Belgium), and introduced new crops in the environment, a factor known to promote virus emergence. In this study, an emergent virus (PhCMoV) was detected in six farms out of 21. Even though the virus was detected in Germany in 2003 (also in a diversified system, Temple et al., 2021), its prevalence in Belgium could potentially be associated with the development of these diversified farms.

Interestingly, on the most diverse farms, the presence of insect-transmitted viruses was not associated with more significant concern among growers or more symptomatic plants, suggesting that their presence was not associated with very high viral disease risk. Some studies postulate that plant-cultivated diversity protects against the spread of viral diseases (Haddad et al., 2009, Keesing et al., 2010, Pagan et al., 2012, Roossinck et al., 2015). Different protection mechanisms are involved. For example, creating a large genetic diversity for plants may also lead to a dilution protective effect since pathogens have more chances of encountering resistant hosts in diverse habitats and spreading difficulties (Liu et al., 2020; Keesing et al., 2021). In addition, plant diversity might increase the diversity of insect-vector but also of the predators, which should theoretically lead to an ecological balance (Cook-Patton et al., 2011, Haddad et al., 2010). It might also disturb the movement of insect’s vector and thus reduce the spread of the disease (Power, 1991). Lichtenberg et al., 2017 demonstrate that both organic farming and higher in-field plant diversity enhanced arthropod abundance, particularly for rare taxa, resulting in increased richness but decreased evenness. Our results align with this statement since insect-transmitted viruses were more present in most diversified production systems but were not associated with higher viral disease risk. Nevertheless, more in-depth studies need to be undertaken to confirm this hypothesis. Alongside its other benefits, plant diversity has been shown to reduce the impact of other pathogens such as fungi on crops and can be used for their management (Mundt et al., 2022, Ratnasass et al., 2010). The agroecological paradigm states that pests and pathogens can be controlled, but not eliminated, through antagonistic ecological processes facilitated by crop genetic diversity or enhanced plant health (Van Bruggen et al., 2016). In addition, diversified farms increase the resilience of the systems to biotic (pest and pathogens), abiotic (climate change) and economic (one product that can no longer be easily sold) threats because if one crop is affected, growers can rely on other crops (Mori et al., 2013, Petersen-Rockney et al., 2021). These systems minimize the negative impact of agriculture on ecosystems and are less dependent on unstable geopolitical contexts, which can hamper the distribution of food from producers to consumers (Wezel et al., 2014, Hertel et al., 2021). Altogether, and in complement to the documented economic potential of diversified production systems (Van der Ploeg et al., 2019), these considerations suggest that additional research efforts on these systems are needed to better optimize them.

Plant viruses constantly evolve, and monitoring is essential to forecast emergence and outbreaks due to viral diseases. In this survey, we highlighted that some detected viruses would require a little more attention than others. For example, PhCMoV was largely distributed, associated with severe symptoms on few plants and considered problematic by one grower. In addition, knowledge of the biology of this virus is limited (Temple et al., 2021); thus, interactions of the virus with plants and vector within the environment (reservoirs plant, transmission…) needs to be investigated more in-depth to understand how the disease can be developed and to set up control strategies. From the limited experience, PhCMoV can be a recurrent problem year after year unless reservoir host plants (not yet fully known) are removed and vector population reduced (Temple et al., 2021).

Other not-well-known viruses were detected in tomatoes, among which two species (CIRV, SLRV) were not detected in tomatoes before. However, since their presence was not associated with specific symptoms when no co-infection was noticed with other pathogenic viruses, and that growers did not notice it, their characterization might not be the priority for the moment.

Eventually, ToMV was detected on symptomatic plants showing typical ToMV symptoms (mosaic) and asymptomatic plants on a farm where they were unaware of tomato viruses. ToMV was considered a severe threat to tomato production worldwide before the use of resistant cultivars and is currently re-emerging with the increased use of older, non-resistant cultivars (Hanssen et al., 2010). The virus is spread by seed and contact, and the grower affected with ToMV re-used its seeds from one year to another. Therefore, more information can be communicated to avoid practices that increase the likelihood of certain viruses spreading.

Finally, some studies suggested that the presence of STV might increase the pathogenicity of other viruses in mixed infection (Gonzalez et al., 2021). Nevertheless, a recent study observed a mutualist interaction between STV (in single infection) and the host plant (Fukuhara et al., 2020). Here, STV was present more frequently in asymptomatic plant pools than in symptomatic pools, suggesting a potential minor role in plant pathogenicity.

## 5 Conclusion

The methodological approach used here makes it possible to obtain a holistic view of issues related to tomato viruses by combining, for the first time, a survey of growes’ perceptions with the characterization of the tomato virome by HTS technologies. In particular, the grower’s perception enabled a more realistic understanding of the impact of the tomato virome on the field and highlighted the need for better characterization of viruses detected by HTS, as at least one can be a threat to production (e.g. PhCMoV). Overall, the results showed that the presence of plant viruses was not necessarily linked to a high disease risk for the production according to the perception of growers and the symptom prevalence. Since more insect-viruses were detected in the most diversified field, without a higher viral disease pressure nor a concern from the growers, our results are in line with the hypothesis that plant diversity might mitigate the impact of plant viruses on crops. Overall, these results are convergent with the hypothesis of Keesing et al., 2010 which postulate that high biodiversity may provide a larger potential source of novel pathogens (virus emergence) but would reduce further transmission for both long established and newly emerging diseases. It would be interesting to develop further research on the role of plant diversity in vegetable farming systems on the emergence and prevalence of plant viruses to evaluate the risks associated with these sustainable productions systems on a long term.

The results also highlight a lack of awareness from small-scale growers in regards virus threat, including ToBRFV, which might be taken in consideration for further communication on quarantine diseases. The part of the study benefit (side effects) was to inform the growers to certain viral threats and their phytosanitary obligations.

This type of research, which links grower perceptions to the presence of viruses detected by HTS, could be applied to many other plants, systems, and contexts to identify potential messages that need to be communicated to growers according to the viruses present on their farms and the territory and their perception. In this study case, we communicated about the lifecycle of the principal important viruses detected in Belgium as growers claimed their lack of knowledge about plant viruses. All the interviewed growers and the extension services were informed about the detected viruses, and links to websites describing the identified viruses were also sent. Such communications and information could be a starting point for farmers increasing awareness of viruses which, in conjunction with support on how to report viruses, and how to safely control the viruses may result in fewer losses from viral disease outbreaks.

## Supporting information

Suppl. data.zip

## 8 Supplementary materials

**Supplementary data 1.** Questionnaire

**Supplementary Figure 1.** Pictures of PhCMoV-infected plants (tomatoes, eggplant, cucumber, galinsoge and capscum) shown to the growers at the Q5 of the questionnaire (Temple et al., 2021)

**Supplementary Figure 2.** Pictures of symptomatic plants, a) Farm: E, Presence CMV, b) Farm: Q, Presence ToMV, c) Farm: S, Presence of PhCMoV

**Supplementary Table 1.** Details of all the gathered characteristics per farm or grower. G: Grower characteristic, F: Farm characteristic, T: Tomato culture characteristic.

**Supplementary Table 2.** List of the different plant viruses detected by farms in symptomatic or asymptomatic pool of plants. The crop and number of symptomatic plants tested by RT-PCR is also indicated. NA= non applicable, + = positive, − = negative

## 10 Author Contributions

CT, AB, SM, and KM contributed to the conception and design of the study and the methodology design. CT, AB, SM, KM, and ST validated the study. CT realized the interviews, field observations, sample collection, and bioinformatics analyzes. CT and SS performed the laboratory analysis. KM provided a strong support for analyzing the interviews. CT wrote the first original draft. CT, AB, SM, KM, ST revised and edited the manuscript. SM and KM provided resources. SM and AB provided supervision. All authors contributed to the manuscript revision, read and approved the submitted version.

## 12 Acknowledgments

**1** The authors would like to thank the National Institute of Biology of Slovenia and especially Denis Kutnjak and Mark Paul Selda Rivarez for their help in bioinformatic analysis. We also acknowledge the team of Phytopathology of Gembloux Agro-biotech, Laurent Minet (Hortiforum asbl / Centre Technique Horticole de Gembloux), Johny Hilaire and the members of the extension services (Centre Interprofessionel Maraicher) for their support and brainstorming effort in designing the survey and the questionnaire. Elisabeth Demonty and Pierre Hellin (Plant Virology Lab, CRA-W) are also thanks for their support in the laboratory analyzes. We are also very grateful to the growers who took the time to reply to the questionnaire and allowed us to access their properties and collect samples. We thank Johan Rollin, François Maclot and Nuria Fontdevila for their support and advice on analyzing the data.

## 14 Ethics statements

Oral consent was given from all the participants before the questionnaire. They were all informed and consented that there was a mandatory notification to the local authorities if a quarantine pathogen was detected in their crops. The university’s ethical committee is mainly focused on medical studies and was therefore unsuitable for evaluating this study.

## 15 Contribution to the field statement (200 words max)

Plant viruses can cause severe diseases in tomatoes, reducing yields and fruit quality. Once a virus has infected a plant, there is no cure. Therefore, viral diseases must be avoided prophylactically. The impact of viruses on tomatoes can vary depending on the biology of each virus, the cultivar and the environment. Diseases caused by viruses account for almost 50% of emerging plant diseases, reinforcing the need of awareness on these pathogens for growers. In highly industrialized countries, the number of small-scale vegetable growers relying on crop diversification and crop rotations has increased recently. However, the risks associated with viruses in these sustainable production systems are unknown. This study aimed to understand the viral disease risks threatening diversified production systems including tomatoes and the impact of cultural practices on these diseases. For this purpose, an innovative methodology was developed, combining for the first-time new technologies for virus identification (high throughput sequencing) and an questionnaire dedicated to the growers. The questionnaire aimed to understand growers’ perception regarding viral diseases’ impact and to describe the typology of the farms, which has been compared with the presence of viruses.

