## Supplementary material for "High Throughput Sequencing technologies complemented by grower’s perception highlight the impact of tomato virome in diversified vegetable farms": Suppl. data.zip: Supplementary_Material-VF.docx

### 1 Supplementary Data

Questionnaire:

#### 1.1 Characterization of the farm (F) and personal background of the growers (P)

What is the size of your farm?

On what area do you grow vegetables?

How many vegetables do you grow on average per year?

How long have you been growing tomatoes on this land?

How long have you been working in vegetable farming?

Have you always worked in agriculture?

If not: what field were you in?

Do you have or did you have family in the agricultural sector?

How many of you work in this structure permanently?

Are you certified organic?

If not: Do you intend to become certified?

What are your marketing channels?

#### 1.2 Technical itinerary of tomato culture: (T)

How many tomato plants do you have?

How many varieties do you grow on average per year?

Where do you get your plants from?

If they sow the seeds: where do you get your seeds from?

Do you reuse your seeds from one year to the next?

If yes: Do you disinfect them before usage?

Do you disinfect your tools regularly?

#### 1.3 Perception on tomato diseases (focus on viral diseases)

See Figure 1

### 2 Supplementary Figures and Tables

#### 2.1 Supplementary Figures

**Supplementary Figure 1.** Pictures of PhCMoV-infected plants (tomatoes, eggplant, cucumber, galinsoge and capsicum) shown to the growers at the Q5 of the questionnaire

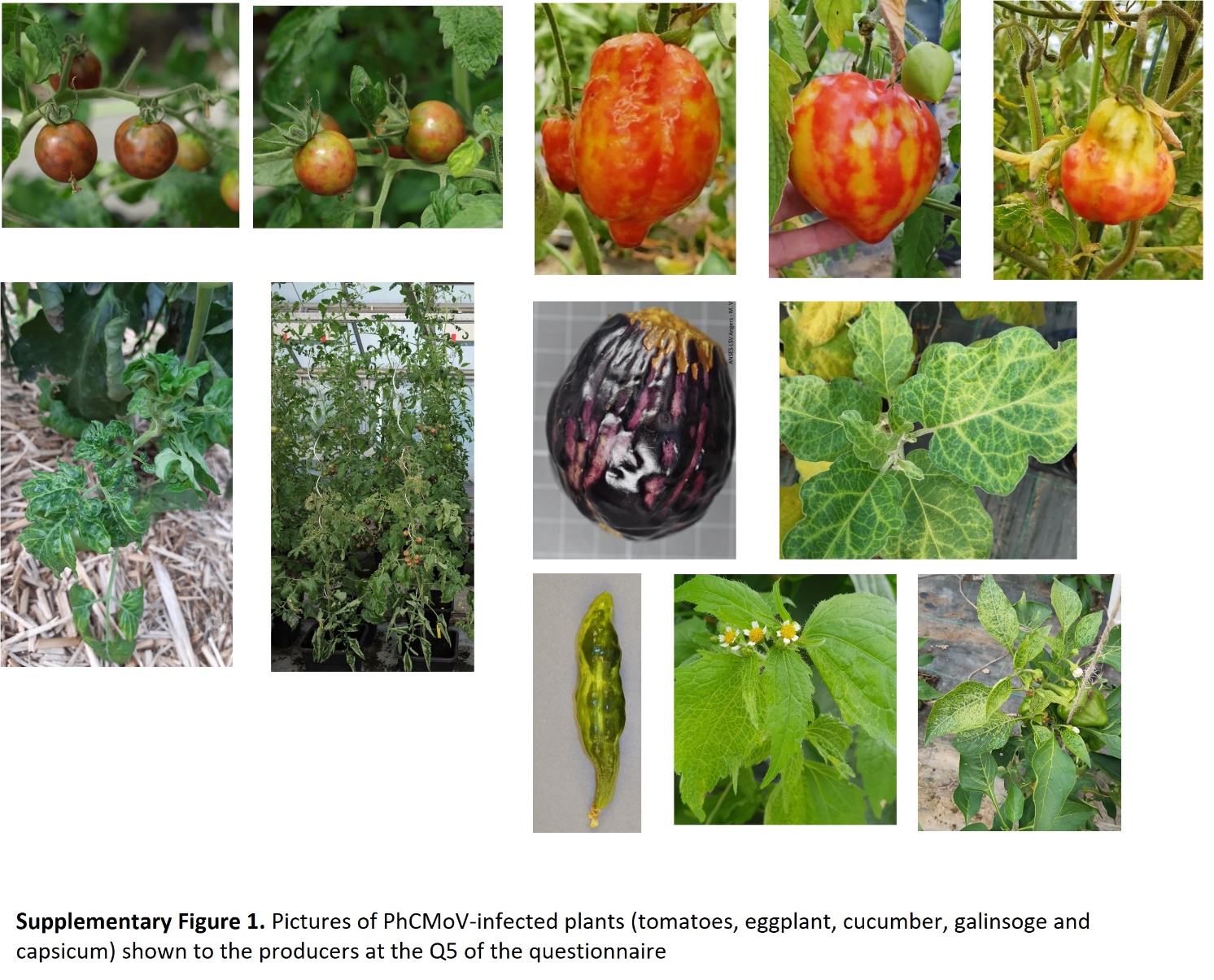

**Supplementary Figure 2.** Pictures of symptomatic plants

a) Farm: E, Presence CMV

b) Farm: Q, Presence ToMV

c) Farm: S, Presence of PhCMoV

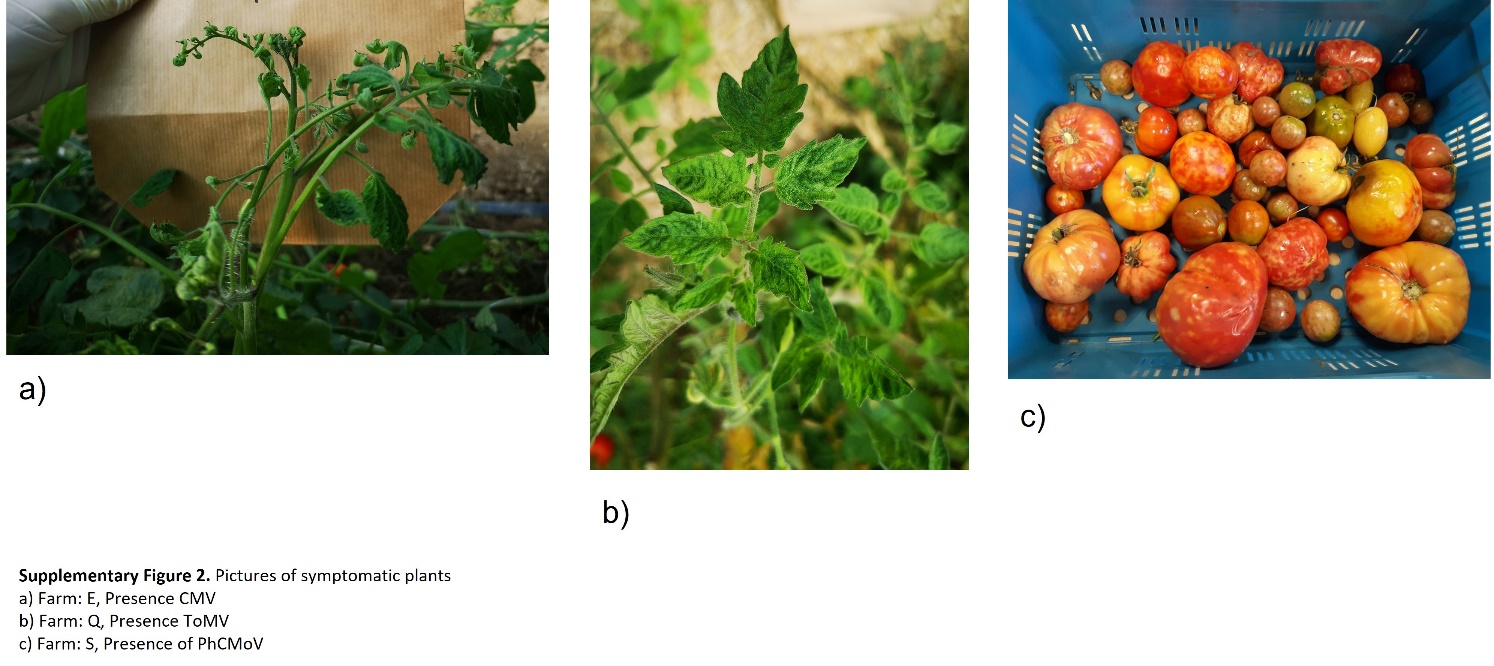

#### 2.2 Supplementary Tables

**Supplementary Table 1.** Details of all the gathered characteristics per farm or grower. G: Grower characteristic, F: Farm characteristic, T: Tomato culture characteristic.

**Supplementary Table 2.** List of the different plant viruses detected by farms in symptomatic or asymptomatic pool of plants. The crop and number of symptomatic plants tested by RT-PCR is also indicated. NA= non applicable, + = positive, - = negative
