## Supplementary figures and images for "High Throughput Sequencing technologies complemented by grower’s perception highlight the impact of tomato virome in diversified vegetable farms"

### Suppl. Fig. 1.jpg

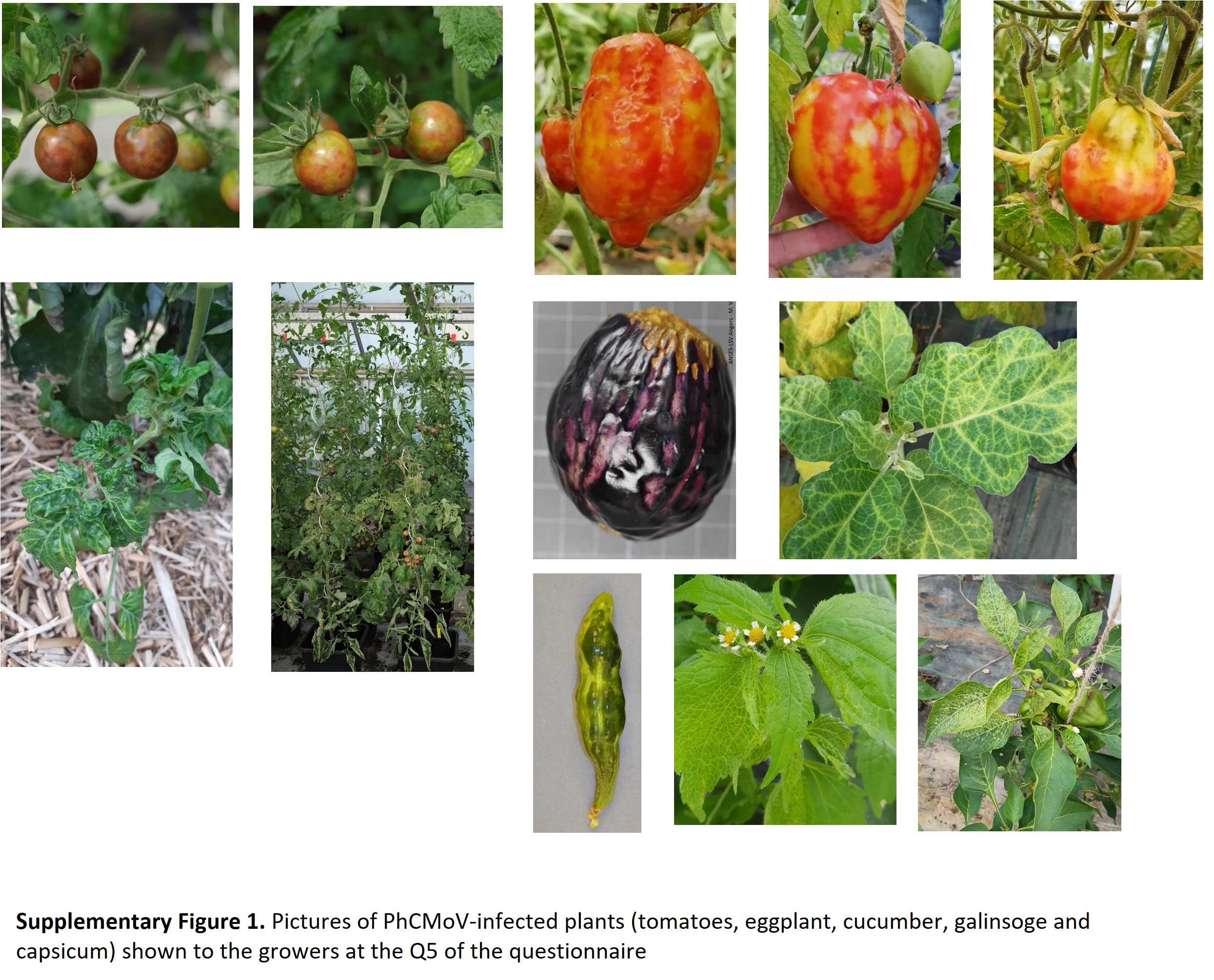

### Suppl. Fig. 2.jpg

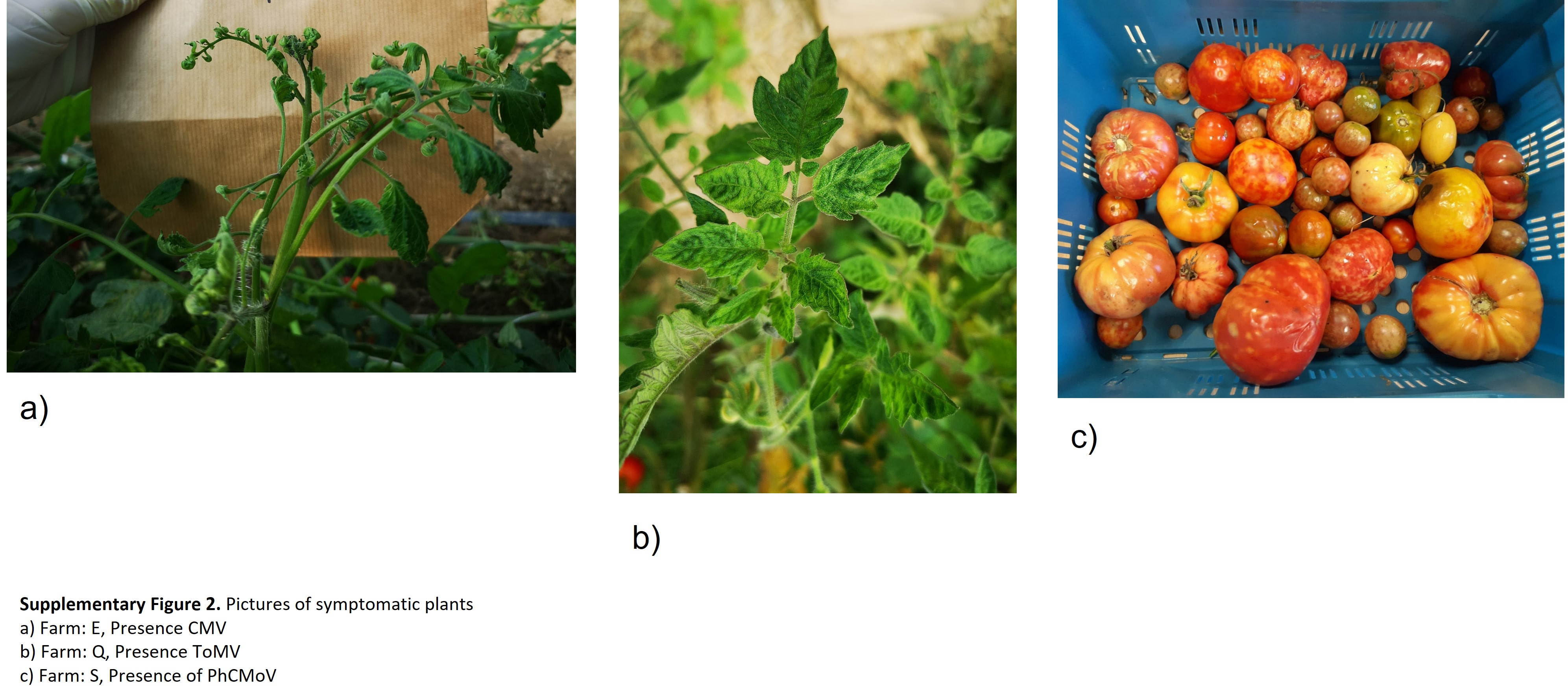
